# Modified tau leads to neuronal and global ribosome dysfunction in *C. elegans* model of tauopathy

**DOI:** 10.1101/2025.08.29.670302

**Authors:** Atalay Ata, Hannah M. Somers, Noah Lind, Jeremy H. Fuqua, Jarod A. Rollins

## Abstract

Alzheimer’s disease (AD) is a neurodegenerative disease characterized by an early loss of memory formation which requires protein synthesis. Tau is an intrinsically disordered protein and is subject to extensive post-translational modifications (PTMs). Some PTMs have been shown to alter localization of tau and allow tau to disrupt protein translation. Protein interactome studies indicate that tau might interact with ribosomal proteins. Therefore, we hypothesized that tau is causing ribosomal dysfunction as an early event and this interaction is dependent on tau’s PTMs. To test this, we used a *C. elegans* strain expressing single copy insertion of human tau as well as two of the most frequent modified versions of tau in mechanosensory neurons. With our assay to measure translation, we showed that in our T231 phosphorylation mimetic strain, there was a significant decrease in neuronal translation. This mimetic strain also showed a significant decrease in median lifespan and locomotion. Unexpectedly, in all our Tau-expressing strains, we detected a significant decrease in whole worm translation, suggesting a possible role of tau to influence translation in other tissues in worm. Our *in vitro*, *in vivo* and *ex vivo* efforts to demonstrate tau-ribosome association via fluorescent polysome profiling have shown that there is no direct association between tau and the ribosome. Ribosome dysfunction caused by modified tau could be an early event in AD pathology before the pathological hallmarks appear.

## Introduction

Alzheimer’s disease (AD) is a neurodegenerative disorder characterized by the progressive loss of memory and cognitive function (Long & Holtzman, 2019). One of the pathological hallmarks of AD is the accumulation of abnormally modified tau protein in the brain, which leads to the formation of neurofibrillary tangles and neuronal death (Guo et al., 2020). Tau pathology is associated with earlier AD onset and accelerated disease progression (Frontzkowski et al., 2022). There is also evidence that Aβ-induced cognitive deficits are dependent on tau in AD mouse models (Ittner et al., 2010). Due to its intrinsically disordered structure, tau protein is subject to extensive post-translational modifications (Alquezar et al., 2021; Mandelkow & Mandelkow, 2012). Some of these modifications are known to alter the binding of tau to microtubules - this can result in disassociation of tau from microtubules and mislocalization to the somatodendritic compartment (Alquezar et al., 2021; Carroll et al., 2021). Specifically, T231 phosphorylation has been shown to be critical for disrupting tau’s binding to microtubules (Schwalbe et al., 2015). Tau K281 acetylation has been shown to affect stabilization of axon initial segment cytoskeleton and contributes to tau’s mislocalization in a mouse model (Sohn et al., 2016). Additionally, tau K274 and K281 acetylations have been shown to contribute to synaptic dysfunction and memory loss related to Alzheimer’s disease (Tracy et al., 2016). Specifically, acetylated tau disrupts postsynaptic signaling required for long term synaptic strengthening (Tracy et al., 2016). In mouse models and patient brains, T231 phosphorylation and K274/K281 acetylation are frequently detected (Wesseling et al., 2020). Accumulation of Tau post-translational modifications are thought to precede aggregation related pathologies in AD (Carroll et al., 2021). Therefore, it is critical to understand the impacts of these modifications on tau function and characterize the potential of modified tau protein to disrupt several cellular processes (Carroll et al., 2021; Dujardin et al., 2020). While the precise mechanisms by which tau pathology contributes to AD pathogenesis are not fully understood, recent studies have implicated an interaction between tau and the ribosome as an early event in the disease’s progression (Kavanagh et al., 2022).

Ribosomes are complex molecular machines that play a central role in protein synthesis by translating mRNA into polypeptide chains. In neurons, ribosomes found in dendrites are involved in the regulation of synaptic plasticity and memory formation (Monday et al., 2022; Sun et al., 2021). Actively translating ribosomes participating in polysomes are upregulated in dendritic spines during memory consolidation (Ostroff & Cain, 2022). Several studies have shown that tau can interact with ribosomal proteins and alter their activity, potentially disrupting the delicate balance of protein synthesis in the brain (Gunawardana et al., 2015; Kavanagh et al., 2022; Tracy et al., 2022; Wang et al., 2019).

Recent work in model systems has provided evidence that an association between tau and the ribosome may be an important early event in AD pathogenesis. Using a human AD and control brain lysates, researchers demonstrated that tau coimmunoprecipitates with ribosomal protein RPS6 and induced expression of Tau decreases the synthesis of ribosomal proteins (Koren et al., 2019). Importantly, these effects can be observed at an early stage of tau pathology suggesting that tau-ribosome association may represent an early molecular event in AD pathogenesis (Koren et al., 2019). Additionally, tau colocalizes with ribosomal proteins in AD patient brain fractions (Meier et al., 2016). Ribosomal proteins are also detected as possible interaction partners of tau in proteomics and quantitative interactome approaches. (Gunawardana et al., 2015; Kavanagh et al., 2022; Tracy et al., 2022; Wang et al., 2019). However, it isn’t known whether tau is interacting with actively translating ribosomes (polysomes – a low-complexity possible early event in the disease pathology. Therefore, we hypothesized that tau and post-translationally modified tau associate with actively translating ribosomes in polysomes to cause ribosomal dysfunction.

To investigate, we utilized *C. elegans* as a model system for neuronal function and longevity due to the availability of techniques to study tau related pathologies (Natale et al., 2020). To this end, we utilized a *C. elegans* strain in which human wild-type tau is expressed in mechanosensory neurons with a single copy insertion system. This strain was further modified to mimic the commonly occurring AD specific post-translational modifications s tau T231 phosphorylation and K274/281 acetylation (Guha et al., 2020). In the original characterization of this stain, investigators assessed the effects of human wild type tau with disease-specific modifications on mechanosensory neuronal function and paraquat induced mitophagy (Guha et al., 2020). In addition, we utilized another pre-existing *C. elegans* strain in which human wild type tau is expressed pan-neuronally (Aquino Nunez et al., 2022). With this strain, investigators assessed the effects of tau and 3PO aggregation prone variant tau accumulation in an age dependent manner (Aquino Nunez et al., 2022). With the strains that we acquired, we wanted to characterize the effect of tau on lifespan and locomotion in the context of neuronal translation. To measure neuronal and global translation levels, we used our modified puromycin assay where we used a modified version of puromycin to label nascently synthesized polypeptide chains in a tissue specific manner and quantify neuronal and global translation (Somers et al., 2022). Lastly, we tried to visualize direct tau-ribosome association with fluorescent polysome profiling (Shaffer & Rollins, 2020).

## Materials and Methods

### Strains

N2 wildtype (Bristol), DC19 bus-5(br19) X, KWN167 rnySi24 [Pmec-7::Dendra2::Tau-T4; unc-119+] II., KWN789 rnySi52 [Tau-T4 (T231E) *rnySi24] II. KWN790 rnySi53 [Tau-T4 (K274Q; K281Q) *rnySi24] II., KWN169 rnySi26 [Pmec-7::Dendra2; unc-119+] II., JAR041 rnySi52 [Tau-T4 (T231E) *rnySi24] II, bus-5(br19) X., JAR040: rnySi24 [Pmec-7::Dendra2::Tau-T4; unc-119+] II, bus-5(br19) X., JAR042: rnySi53 [Tau-T4 (K274Q; K281Q) *rnySi24] II, bus-5(br19) X., JAR043: rnySi26 [Pmec-7::Dendra2; unc-119+] II, bus-5(br19) X., JAR020 rps-6(rns6[rps-6::mCherry]); juSi123 [rpl-29::GFP] II., NW1229 evIs111 [Prgef-1::GFP], EVL1655 lhIs95 [Prgef-1::hTau40::GFP]. KWN strains were provided by Dr. Keith Nehrke at University of Rochester. NW1229 and EVL1655 were provided by Dr. Brian Ackley at University of Kansas. The DC19 bus-5(br19) mutant was obtained from the CGC. Worms were grown at 20°C on nematode growth media (NGM) agar plates seeded with OP50 bacteria at 2 x 10^10 CFU. Hermaphrodites were utilized for all assays. To age-synchronize populations, adult worms were bleached unless mentioned otherwise.

### Fluorescent polysome profiling in *C. elegans*

Fluorescent polysome profiling was performed as described (Shaffer et al.). Briefly, 10,000 to 20,000 age synchronized day 1 adult worms were pelleted, and 100 ul of pelleted worms were lysed with 350 ul high-salt solubilization buffer (300 mM NaCl, 50 mM Tris–HCl [pH 8.0], 10 mM MgCl2,1mM EGTA, 400 U RNAsin/mL, 0.2 mg cycloheximide/mL, 1% Triton X-100, and 0.1% sodium deoxycholate). After solubilization, lysates were incubated with an additional 200 ul of solubilization buffer in a rotating incubator at 4°C for 30 minutes. The lysates were centrifuged at 14,000 g for 5 minutes to pellet debris. The supernatant was measured for RNA at 260 nm absorbance for equalizing the load on sucrose gradient. After determining A260, samples were loaded on 5% to 50% sucrose gradient in high salt resolving buffer (140 mM NaCl, 25 mM Tris–HCl [pH 8.0], and 10 mM MgCl2) and centrifuged in a Sorvall TH-641 rotor at 38,000 rpm for 2 h @ 4 °C. Gradients were fractionated using a Piston Gradient Station equipped with a dual fluorescent channel capable Triax flow cell (BioComp Instruments) with continuous monitoring of absorbance at 260 nm and peak fluorescence emission at 470 nm with a bandpass of 40 for GFP and peak fluorescence emission at 645 nm with a bandpass of 75 for mCherry.

### O-propargyl-puromycin (OPP) assay for quantifying neuronal and global translation in *C. elegans*

O-propargyl-puromycin assay was performed as described (Somers et al.). Briefly, 100 day one adult worms were harvested from OP50 seeded NGM plates by washing with liquid NGM into a 1.5 mL microcentifuge tube. Additional OP50 was removed via brief washes in liquid NGM. Worms were incubated for 3 hours at 20 °C with shaking in 10 µM O-propargyl-puromycin diluted in liquid NGM containing 2 x 10^8 CFU/mL of PFA killed OP50 in a 1.5 mL microcentrifuge tube. Worms were fixed with 100 µl of 4 g of paraformaldehyde in RNase-free PBS. After washes, conjugation of fluorophore to puromycylated peptides was carried out according to the Click-iT® Plus Alexa 647 Fluor® Picolyl Azide Tool Kit. Fixed worms were incubated in Click-iT reaction buffer overnight at 10°C. Worm pellets were washed with PBS three times before imaging.

3D images for the OP-Puromycin assay were collected with a Zeiss LSM 980 confocal microscope (Carl Zeiss Microscopy) on an upright Zeiss Axio Examiner stand equipped with a Zeiss Plan Apo 20X/0.8NA objective (Carl Zeiss Microscopy). DAPI fluorescence was excited with the 405 nm line with 1% laser intensity, from a 30-mW laser diode (Laser beam attenuation via direct modulation). EGFP fluorescence was excited with the 488 nm line, 8% laser intensity from a 30-mW laser diode for mechanosensory strains and 1% laser intensity for the pan-neuronal strains. Alexa 647 fluorescence was excited with a 639 nm line, 1% laser intensity from a 25-mW laser diode. Laser beam attenuation for all channels other than DAPI was by using visible range acousto-optic tunable filters. Images were captured using Airyscan 2 with the following detection wavelengths: DAPI from 422 to 497 nm, Dendra2 from 493 to 517 nm and 659 to 720 nm, EGFP from 499 to 557 nm and 659 to 720 nm, Alexa 647 from 499 to 557 nm and 659 to 720 nm. Images were sequentially acquired in Super Resolution mode (SR) at zoom 1.7, with a line average of 1, a resolution of 512 3 512 pixels, a pixel time of 0.85 us, in 8-bit and in bidirectional mode. Z-stack images were collected with a step size of 0.31 mm with the Motorized Scanning Stage 130 3 85 PIEZO (Carl Zeiss Microscopy) mounted on Z-piezo stage insert WSB 500 (Carl Zeiss Microscopy).

Microscope was controlled using Zen Blue software (Zen Pro 3.1), Airyscan images were processed manually in 3D with a strength of the deconvolution set to 4.1 and saved in CZI file format. All imaging parameters were identical between experiments for compared images. Settings were different between pan-neuronal and mechanosensory OPP imaging. The image analyses were done on the visible mechanosensory neuron somas or all somas as well as entire worm body by using Imaris v.9.5. Raw data were segmented by manually outlining the mechanosensory neurons or all neurons using the *mec7::Dendra2* or *Prgef-1::GFP* signal by navigation through the Z-stack. The segmented rendered surface was used to mask the raw data and create a subset on which OPP-Alexa 647 and Dendra2 or GFP intensity measurements were performed on the selected mechanosensory somas or all somas using the surfaces statistics tool for each sample. The automatic Imaris detection was first validated manually, by examining individual z-planes using the Oblique slicer tool and a combination of 2 Clipping plane tools to isolate and examine neuronal specific tissue region. Threshold settings were manually adjusted for each sample to yield the best possible detection.

### Lifespan and locomotion analysis with NemaLife microfluidic chamber in *C. elegans*

NemaLife lifespan assay was performed as described with modifications (Rahman et al.). Around 70 age synchronized day 2 adult worms were loaded on NemaLife microfluidic chambers to be monitored throughout their lifespan with daily wash/feed cycles and locomotion tracking. The worms were free to move in liquid NGM (NaCl, Peptone, 5 mg/mL cholesterol in ethanol, 1 M KPO4 buffer pH 6.0, 1 M MgSO4, 1 M CaCl2) and fed daily with PFA killed E. coli OP50 at 5x10^9 CFU/mL suspension in liquid NGM. The NemaLife Infinity processor allowed us to stimulate the movement of worms and dispose of progeny by washing them with fresh liquid NGM every day and subsequent tracking of the movement. Wash/feed cycle consists of 30 seconds of recording before the wash flow, 30 seconds of washing (1 minute total) and immediately after the wash flow; tracking for locomotion for 90 seconds at 5 frames per second with the shutter speed of 1/20 and focal ratio of 24. After tracking for 21 days, the video files were processed for background subtraction to help distinguish live worms from dead worms on a custom FIJI script. Then, the tracking videos are manually analyzed on WormLab version 2022.1.1 software to generate lifespan and locomotion data. Each day, the number of worms was counted to generate the survival curve and their movement across 150 frames (corresponds to 30 seconds or 1/3^rd^ of the recorded movement) were tracked by the software to measure maximum speed.

### *C. elegans* development assay

Approximately 20 worms were plated to 60 mm NGM plates for 1 hour of egg laying (1 hour incubation at 20°C). 10 of the laid eggs per independent replicate were transferred to individual 35 mm NGM plates at the L1 stage, development was recorded as the time taken for the first egg to be laid as checked hourly.

### *In vitro* recombinant protein labeling

We used a carrier-free human recombinant 0N4R tau protein (R&D Systems, Cat# SP-499) to be labeled with Alexa Fluor™ 594 Microscale Protein Labeling Kit (Thermo Fisher, Cat# A30008) following the manufacturers instruction. Briefly, we calculated an approximation of number of ribosomes per worm and extrapolated that to the number of worms required for our profiling experiments (Raveh et al., 2016). Given the labeling reaction kit equation, [(ug protein/protein MW)*1000]*protein molar ratio/12.2= ul of dye, we calculated 2.78 ul dye required to label 40 ug of protein needed for the 1to1 tau-ribosome association (The required amount to saturate ribosomes was 6.38 ug). 40 kDa is the predicted weight of the purchased recombinant tau protein and protein molar ratio (MR) for a 40 kDa protein is 34 for optimal labeling (Molar ratio is taken from the manufacturer’s instruction). After the room temperature incubation for 15 minutes for the labeling reaction, we proceed to determine the concentration of the labeled protein with A280 and A590 measurements. After determining the concentration, we proceed into mixing 10 ug of labeled recombinant tau protein with N2 supernatant and incubated it for 1 hour at room temperature, which was followed by fluorescent polysome profiling.

### *Ex vivo* fluorescent polysome profiling of frozen mouse cerebella

We acquired frozen mouse cerebellar tissue from Dr. George Canet and Prof. Dr. Emmanuel Planel at Department of Psychiatry and Neuroscience, Faculty of Medicine, Université Laval in Canada (Guisle et al., 2022). The frozen scrap cerebellar tissue is from the B6.Cg-Mapttm1(EGFP)Klt-Tg(MAPT)8cPdav/J strain where EGFP tagged human Tau is expressed in all neurons. We processed 40-210 mg of frozen tissue with 5 mm 440C stainless steel grinding ball (OPS Diagnostics, Cat# GBSS 196-2500-10) in 2ml round bottom tubes in a QIAGEN TissueLyser II instrument with 20 Hz frequency for 1 minute in high-salt solubilization buffer (300 mM NaCl, 50 mM Tris–HCl [pH 8.0], 10 mM MgCl2,1mM EGTA, 400 U RNAsin/mL, 0.2 mg cycloheximide/mL, 1% Triton X-100, and 0.1% sodium deoxycholate) and low-salt solubilization buffer (50 mM KCl, 50 mM Tris–HCl [pH 7.4], 10 mM MgCl2,1mM EGTA, 400 U RNAsin/mL, 0.2 mg cycloheximide/mL) without detergents for fluorescent polysome profiling.

### Statistical analysis

Statistical analyses for lifespan and locomotion data were performed by R version 4.1.2 and RStudio version (2022.02.0) with ggpubr package. Statistical analyses for O-propargyl-puromycin (OPP) assay were performed by Prism 9.2.0 with alpha error level of p < 0.05 considered to be significant. OPP assay data were averaged and represented as mean ± SD. Normalization tests were applied before assessing group differences. Differences in lifespan were obtained by log rank analysis using the survminer package.

### Western Blotting against tau in polysomal fractions

For western blotting, 30 mg of total protein was separated on 4-20% mini-Protean TGX stain-free gels (Bio-Rad). The resulting gels were exposed to UV for 5 min to allow fluorescent detection of total protein present before electroblotting samples to PVDF. Blots were then imaged under UV to image total protein transferred to them. Anti-tau antibody (Cat# MAB3420 Fisher Scientific) was diluted 1:2000 in Everyblot (Bio-Rad) and incubated overnight at 4°C with shaking. Another Anti-tau antibody (Cat# AHB0042 Thermo Fisher) was diluted 1:500 in Everyblot (Bio-Rad) incubated overnight at 4°C with shaking. Anti-phospho Tau T231 antibody (Cat# ab151559 Abcam) was diluted 1:1000 in Everyblot (Bio-Rad) and incubated at RT for 4 h with shaking. After washing, the blot was incubated with Peroxidase AffiniPure Goat Anti-Mouse (Cat# 115-035-044 Jackson Immuno Research) 1:5,000 in Everyblot. Three washes of Everyblot were used after each incubation step. Clarity Western ECL kit was used for chemiluminescence according to the manufacturer’s instruction (Bio-Rad, Cat#1705061). Images of total protein and of chemiluminescence were background subtracted and quantified using ImageJ.

## Results

### Lifespan and locomotion of *C. elegans* models of tauopathy

To characterize the mechanosensory tau strains and pan-neuronal tau strain, we performed lifespan and locomotion analysis with NemaLife microfluidic chamber system. With NemaLife system, we were able to capture the entire lifespan of the strains during which locomotion was assessed with daily wash/feed cycle (Rahman et al.). Our results showed that in mechanosensory tau strains, T231 phosphorylation mimetic tau and K274/K281 acetylation mimetic tau strains had 12.5% and 6.2% decrease in median lifespan compared to N2 control strain respectively (**Fig. 1A**). However, there was no significant difference in lifespan in the pan-neuronal tau strain compared to pan-neuronal GFP control, suggesting a PTM dependent effect of tau (**Fig. 1C**).

**Fig. 1.**
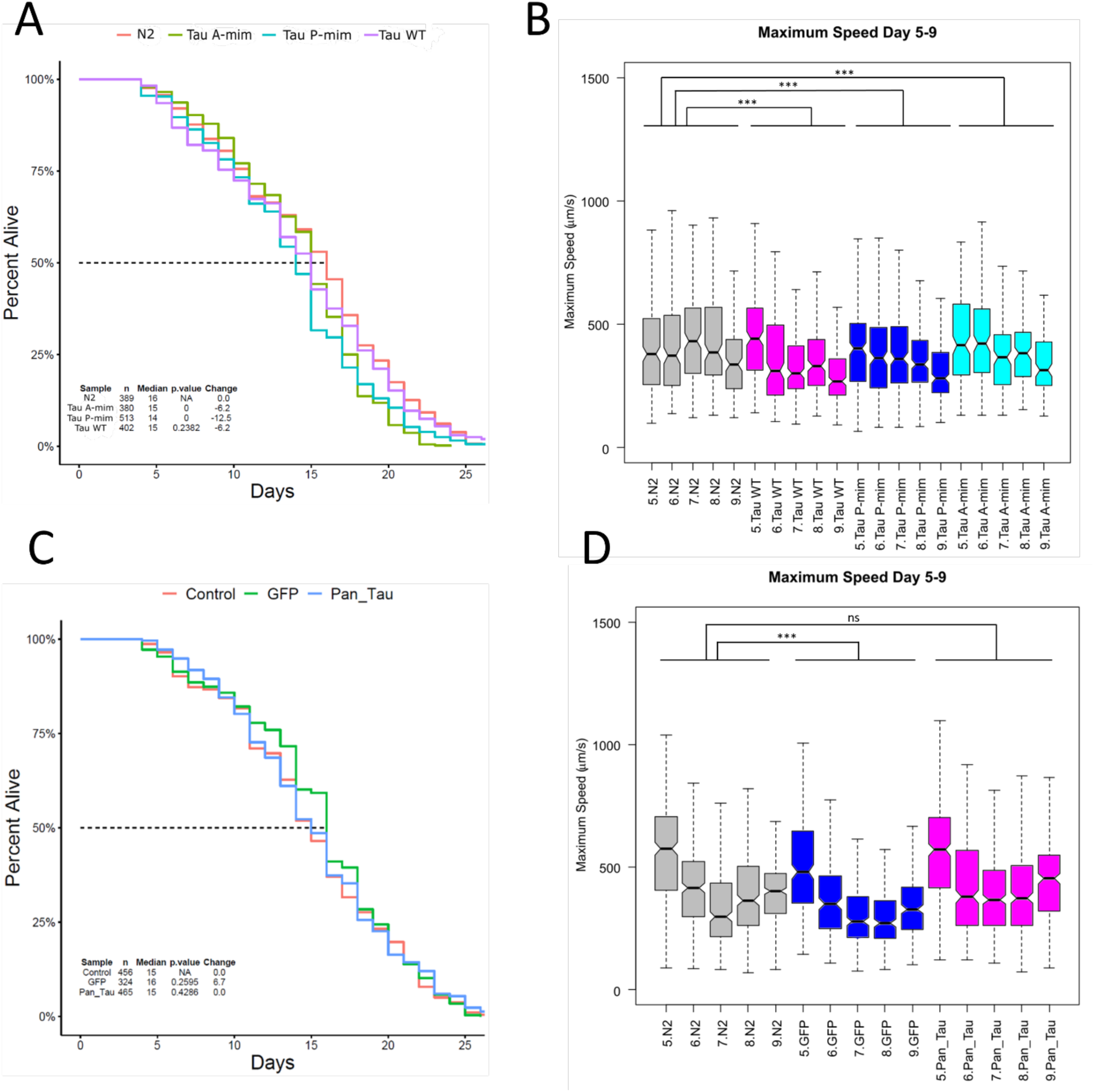
Lifespan and maximum speed of mechanosensory and pan-neuronal tau strains between days 5-9 of adulthood. **A** Tau P-mim strain and Tau A-mim strain have a 12.5% and 6.2% significant decrease in median lifespan, respectively. Tau WT: Wild type tau, Tau P-mim: Tau T231 phosphorylation mimetic, Tau A-mim: Tau K274/K281 acetylation mimetic **B** Wild type Tau, Tau P-mim and Tau A-mim strains had 16.6%, 13.6%, 5.7% significant decrease in maximum speed between days 5-9 of adulthood respectively. Gray: N2, Magenta: Wild type tau, Blue: Tau P-mim, Cyan: Tau A-mim. **C** Pan-neuronal wild type Tau strain didn’t have significant change in median lifespan. GFP: Prgef-1::GFP, Pan_Tau: rgef-1::hTau40::GFP. Statistical analysis was done by one-way ANOVA followed by Tukey’s honest significance test with p value 0 denoting significance and N as the total number of worms across three independent replicates. Dotted line denotes median lifespan. **D** Pan-neuronal wild type Tau strains didn’t have significant change in maximum speed between days 5-9 of adulthood. Gray: N2, Blue: GFP control strain, Magenta: Pan_Tau. Statistical analysis was done by one-way ANOVA followed by Tukey’s honest significance test with ***p < 0.001 denoting significance across three independent replicates. Notches in box plots indicate the estimated 95% confidence interval of the median value (black line). n ≥ 324 aggregated across three independent biological replicates.

The maximum movement speed of tracked animals was compared between days 5 and 9 of adulthood, a period of time where wild type animals typically suffer significant losses in movement speed (Rollins et al 2017). In maximum speed of mechanosensory tau strains between days 5 and 9, wild type tau, T231 phosphorylation mimetic tau and K274/K281 acetylation mimetic tau strains had a 16.6%, 13.6%, 5.7% significant decrease compared to the control strain respectively (**Fig. 1B**). The effect on movement was more pronounced and consistent across multiple days compared to the mild lifespan impairment suggesting a role of tau in altering body wall muscle related mechanisms. In maximum speed of pan-neuronal strains between days 5 and 9, there was no significant difference between pan-neuronal wild type tau and control strain (**Fig. 1D**). Wild type, T231 phosphorylation mimetic tau and K274/K281 acetylation mimetic tau strains resulted in a significant decrease in maximum speed in early adulthood in mechanosensory tau strains compared to wild type tau in pan-neuronal strain. Since we noticed impairments in early days of adulthood, we directed our attention to look for changes in translation levels in early adulthood.

### Quantification of neuronal translation in *C. elegans* model of tauopathy

Ribosomal proteins have been identified as binding partners of Tau in interactome studies (Kavanagh et al., 2022) along with reports that translation is decreased in murine cell culture in the presence of tau (Evans et al., 2021; Meier et al., 2016). Therefore, we wanted to determine if a decrease in protein translation occurred in our *C. elegans* models of Tauopathy as an underlying mechanism for the observed effects on health span and lifespan by Tau. Puromycin based techniques have been developed in *C.elegans* to measure translation tissue specifically (Somers et al., 2022). In this study, we quantified OPP incorporation in neurons as well as whole animals. (**Fig. 2**). In our mechanosensory tau strains, only tau T231 phosphorylation mimetic caused a 39% significant decrease in OPP incorporation in mechanosensory somas compared to controls **(Fig. 2A**). We measured the signal coming from mechanosensory somas and neurites combined from the six mechanosensory neurons in *C. elegans* namely ALML/R, PLML/R and AVM/PVM (**Fig. 2E-H**). The detection and imaging of mechanosensory neurites in their entirety was difficult due to the nature of the Dendra2 fluorophore and we chose to quantify OPP fluorescence in mechanosensory somas for consistency. The quantification was performed from the aggregated somas per worm imaged. The 39% decrease in neuronal translation in mechanosensory tau was not replicated in a pan-neuronal strain where wild type 2N4R tau was expressed under *rgef-1* promoter (**Fig. 2C**).

**Fig. 2.**
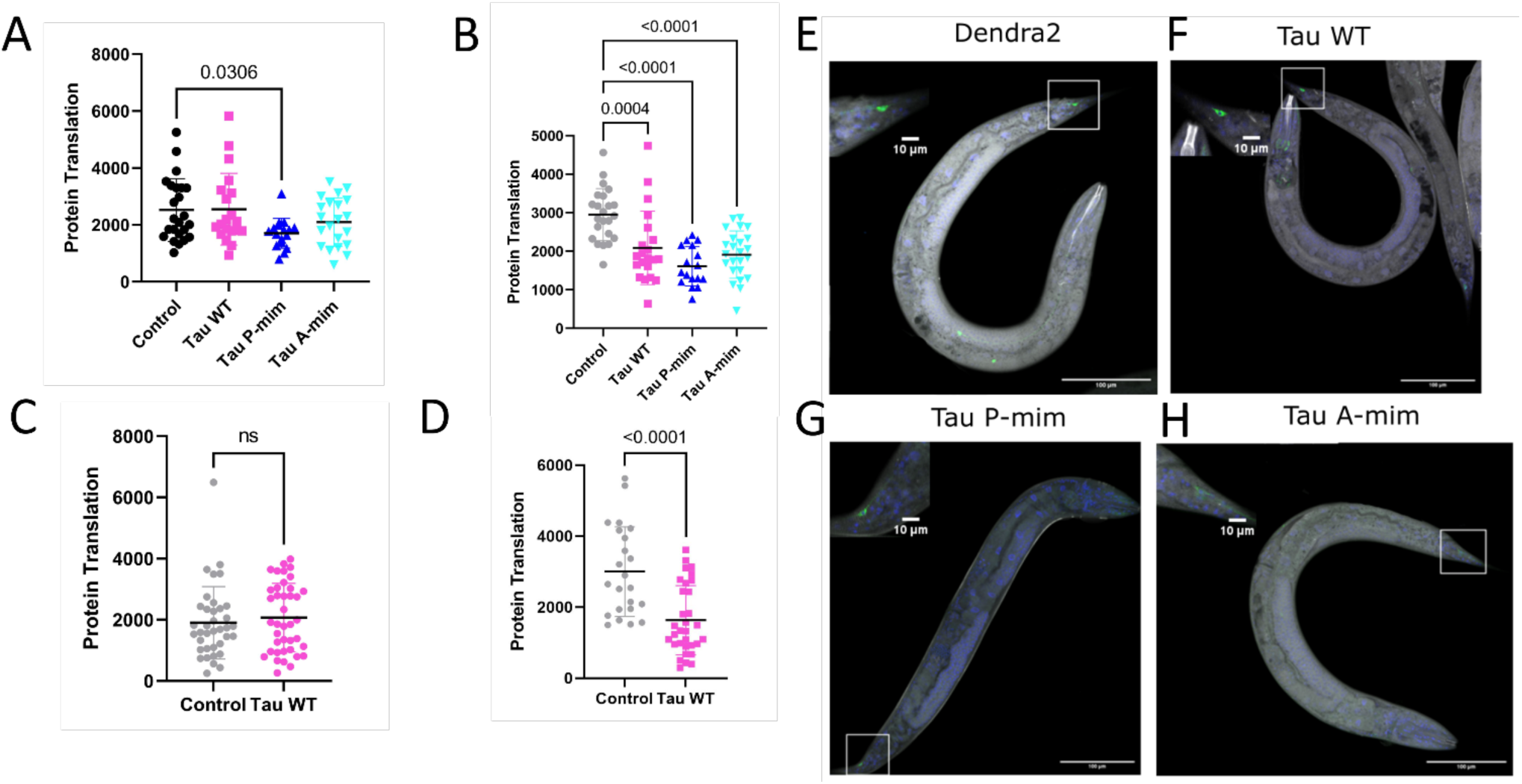
Quantification of OPP Fluorescence that is proportional to protein translation in neurons and whole worms. **A** OPP Fluorescence in mechanosensory somas (N: >20, ANOVA with Dunnett’s m.c.). **B** OPP fluorescence from mechanosensory strain whole worms (N: >20, ANOVA with Dunnett’s m.c. F(3, 81) = 13.82). **C** OPP fluorescence in pan-neuronal Tau strain from all somas (N: >22, Mann-Whitney). **D** OPP fluorescence in pan neuronal Tau strain from whole worms (N: >22, T-test). Tau P-mim strain had 39% significant decrease in neuronal translation in day 1 adult worms. Pan neuronal wild type Tau strain didn’t have a significant decrease in neuronal translation in day 1 adult worms. All strains showed a significant decrease in whole worm translation. All data points are expressed as mean ± SD. Tau_wt: Wild type tau, Tau_Phos: Tau T231 phosphorylation mimetic, Tau_Ace: Tau K274/K281 acetylation mimetic, GFP: Prgef-1::GFP, Pan_Tau: rgef-1::hTau40::GFP. Representative image of Tau mutant strains from O-propargyl-puromycin assay. **E** Inset shows OPP fluorescence (gray) masked over Dendra2 (green) expressed in PLML/R mechanosensory somas. **F** Inset shows OPP fluorescence (grayscale) masked over Dendra2 (green) tagged wild type Tau expressed in PLML/R mechanosensory somas. **G** Inset shows OPP fluorescence (gray) masked over Dendra2 (green) tagged T231 phosphorylation mimetic Tau expressed in PLML/R mechanosensory somas. **H** Inset shows OPP fluorescence (gray) masked over Dendra2 (green) tagged K274/K281 acetylation mimetic Tau expressed in PLML/R mechanosensory somas. Scale bars are 100 µm and 10 µm.

All strains including mechanosensory and pan-neuronal tau showed a striking phenotype in that global OPP signal - which is the entire OPP incorporation in all tissues except neurons - was significantly decreased compared to the control strain (**Fig. 2B, 2D**).

### Resolving direct tau-ribosome association with fluorescent polysome profiling

We initially hypothesized that tau might be interacting with actively translating ribosomes to cause ribosomal dysfunction. After our measurements with respect to neuronal translation, we wanted to visualize the hypothesized direct tau-ribosome association via fluorescent polysome profiling. From our fluorescent polysome profiling experiments, we showed that there was no association between tau, monosomes and polysomes (**Fig. 3**). This was observed for mechanosensory and pan-neuronal tau strains in our low-salt fluorescent polysome profiling conditions (Pringle et al., 2019; Shaffer & Rollins, 2020). In our repeated runs with our tau strains, we were not able to resolve direct association of tau with polysomes compared to a protein that is not a core ribosomal subunit protein such as *rack-1* (**Fig. 3F**).

**Fig. 3.**
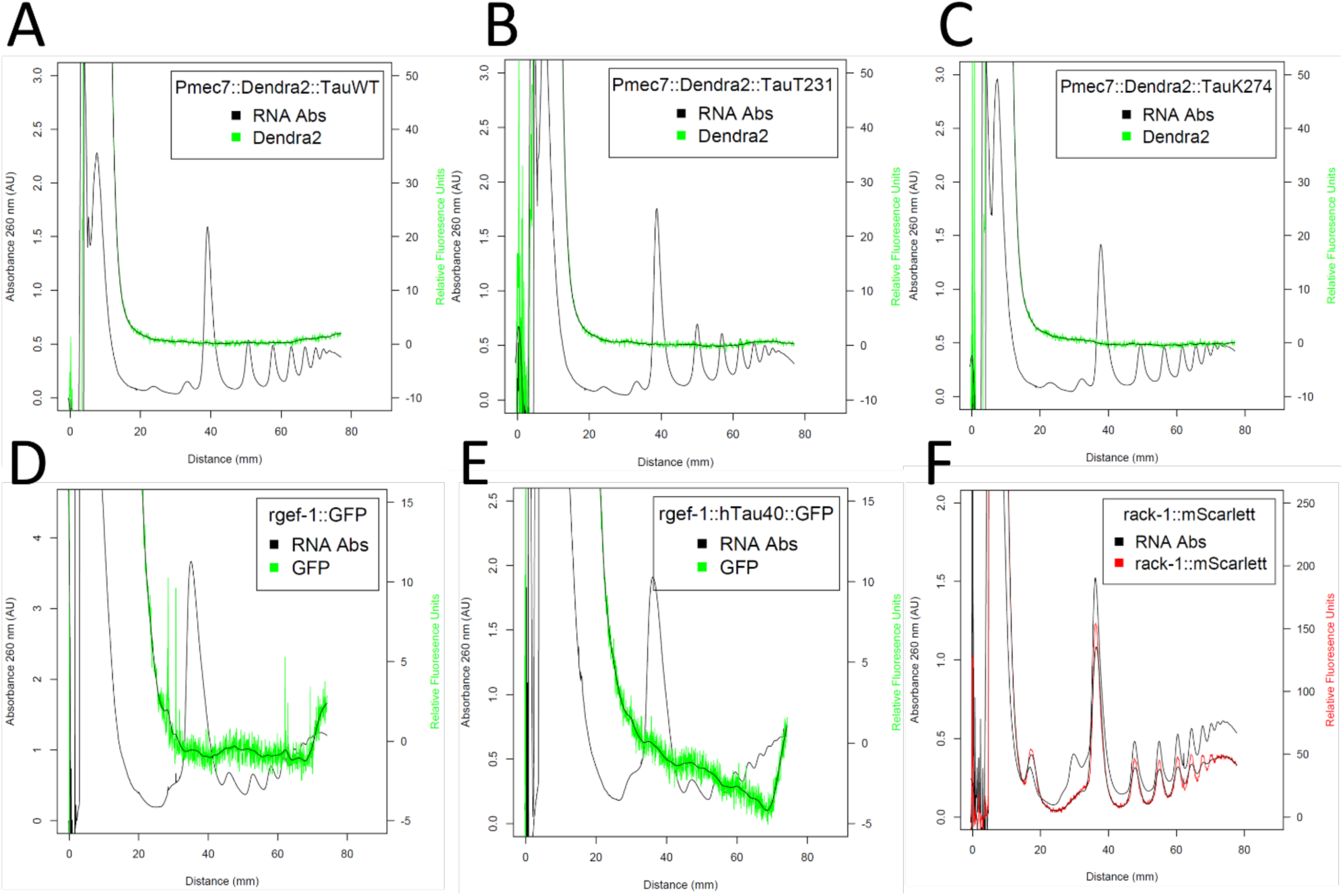
Representative fluorescent polysome profiles of mechanosensory and pan-neuronal Tau strains showing no association with polysomes. **A** Dendra2 tagged wild type Tau strain showing no association with polysomes. **B** Dendra2 tagged Tau T231 phosphorylation mimetic strain showing no association with polysomes. **C** Dendra2 tagged Tau K274/K281 acetylation mimetic strain showing no association with polysomes. **D** Pan-neuronal control strain where GFP is driven by *rgef-1* promoter showing no association with polysomes. **E** GFP tagged wild type Tau expressed pan-neuronally showing no association with polysomes. **F** mScarlet tagged endogenous *rack-1*, which is not a core ribosomal subunit protein showing association from large subunit, monosomes and polysomes.

After our observation in *C. elegans* strains that there was no association, we sought to have an *in vitro* strategy where we acquired a human recombinant tau protein and used a microprotein labeling kit to conjugate the recombinant protein to a fluorophore and mix it with N2 lysate to saturate the ribosomes to visualize association. After three independent replicates, we were able to confirm that the conjugated human recombinant tau also showed no association with polysomes (**Fig. 4**). We also attempted an *ex vivo* strategy where we acquired frozen mouse cerebella where EGFP tagged tau is expressed (Guisle et al., 2022). After two replicates, we were able to show that the EGFP tagged tau in frozen mouse cerebellum showed no association with polysomes **(Fig. 4D**). Lastly, we measured nucleolus area in PLML/R neurons in our mechanosensory tau strains to possibly explain our translational downregulation (Maina, Bailey, Doherty, et al., 2018; Maina, Bailey, Wagih, et al., 2018). We detected a significant decrease in the nucleolus area in PLML/R neurons in mechanosensory tau strains across independent replicates (**Fig. 4F**).

**Fig. 4.**
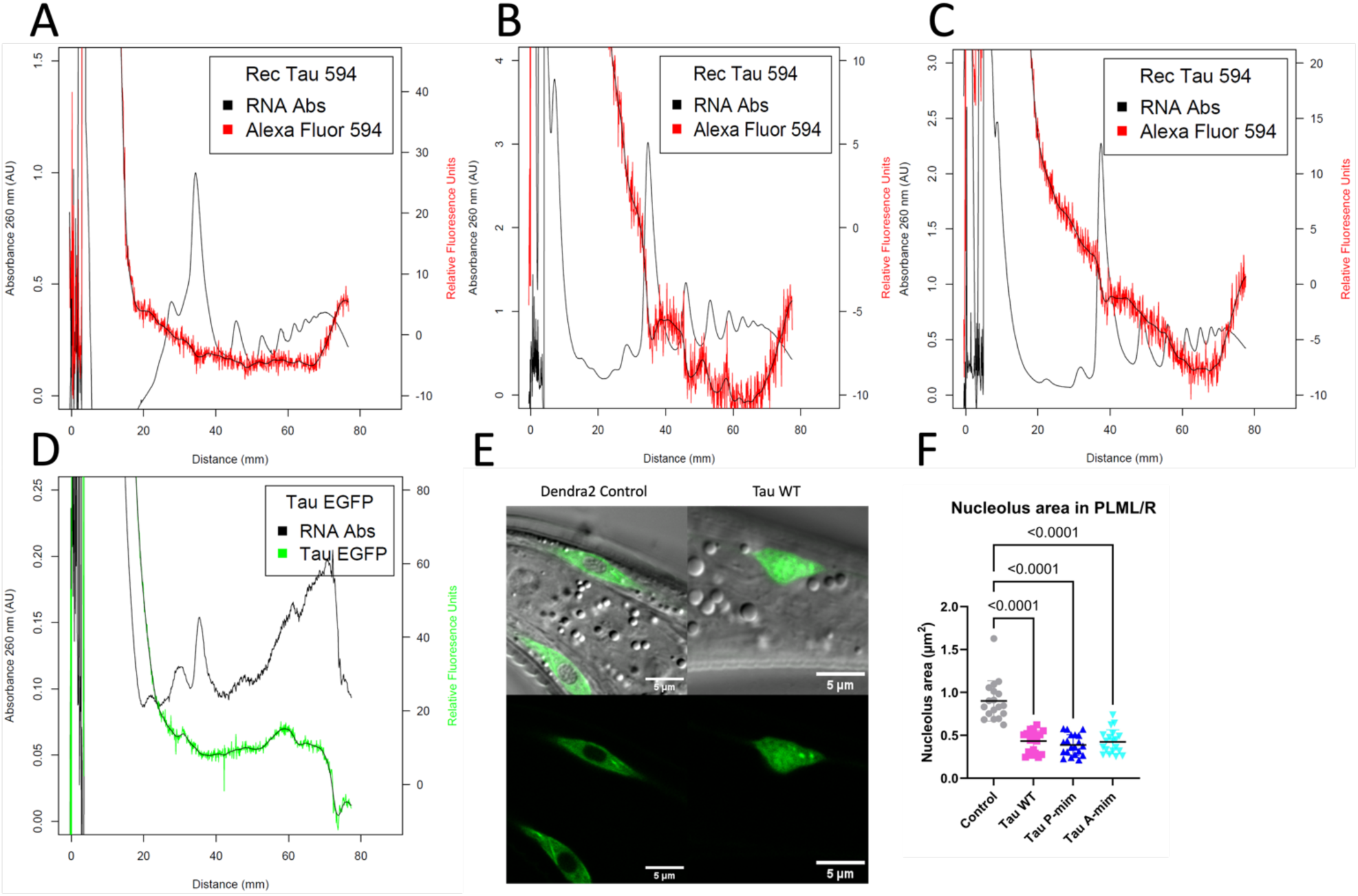
Representative fluorescent polysome profiles of Alexa Fluor 594 conjugated human recombinant tau protein mixed with N2 lysate showing no association with polysomes. **A** Independent biological replicate with an unknown amount of conjugated protein mixed and run for profiling showing no association with polysomes. **B** 10 µg of conjugated protein mixed and run for profiling showing no association with polysomes. **C** 10 µg of conjugated protein mixed and run for profiling showing no association with polysomes. **D** Representative fluorescent polysome profiling of EGFP tagged tau in frozen mouse cerebella showing no association with polysomes. **E** PLML/R nucleoli imaging in mechanosensory tau strains showing nucleolar tau signal. **F** Quantification of nucleolus area in PLML/R neurons showing significant decrease in all mechanosensory tau strains.

## Discussion

We sought to identify tau-ribosome association as an early event in tauopathy and Alzheimer’s disease. To test this, we acquired strains where human wild type tau is expressed either in six mechanosensory neurons or all neurons in *C. elegans*. When we subjected these strains to our microfluidic chamber analysis for lifespan and locomotion, we observed a significant impairment in mechanosensory tau strains suggests that tau expressed in these strains over time has a deleterious health effect and impairs movement (**Fig. 1A**). Since there’s a common pattern in age related changes in locomotion impairment stretching from *C. elegans* to humans, we were able to recapitulate this with our strains (Marck et al., 2016).

The decrease in maximum speed of the worms was more consistent across multiple days of young to middle adulthood (**Fig. 1B**). Assuming a cell non-autonomous role of tau to influence decrease in translation in other tissues, it could be stated that over time this leads to impairments in locomotion. In pan-neuronal strain, there was no significant difference in either lifespan or locomotion suggesting a PTM dependent effect in mechanosensory tau strains (**Fig. 1C,D**). It is known that tau modifications increase the disassociation of tau from microtubules and allow it to relocalize and disrupt several processes (Carroll et al., 2021; Sohn et al., 2016; Tracy et al., 2016). Considering the mild but significant lifespan impairment as well as the locomotion (**Fig. 1A, 1B**), we conclude that the disease specific modifications of tau exacerbate pathology in these strains. Although, in pan-neuronal tau strains, tau is expressed in all neurons compared to mechanosensory neurons, the expected movement phenotype was not observed. In this case, we could not rule out that there could be isoform dependent effects determining tau’s interactions due to the fact that in these strains 2N4R tau is expressed (Aquino Nunez et al., 2022). It is known that isoform differences in tau have an effect on tau’s expression and truncation activity (Boyarko & Hook, 2021).

Our OPP assay showed that in day 1 adult worms, confined to the tau T231 phosphorylation mimetic strain, there was a significant decrease in neuronal translation (**Fig. 2A**). Tau is reported to sequester an eukaryotic ribosome and leading to tau-rRNA aggregates in a yeast system (Banerjee et al., 2020). We know that neurons have a dynamic ribosome system where they can exchange certain ribosomal proteins on their translating ribosomes in a context dependent manner and several CNS pathologies show translation impairments (Fusco et al., 2021; Gamarra et al., 2021). Neuronal activity and local translation are tightly coupled and have an important role in learning and memory. There is a known mechanism in dendrites where during stimulation, a non-canonical translation initiation factor is recruited and translational machinery is rapidly reprogrammed (Hacisuleyman et al., 2024). Therefore, tau compromising neuronal translation as early as day 1 could be a novel mechanism to curb neuronal homeostasis. This is more apparent given the view that over 75% of the both excitatory and inhibitory synapses have protein translation machinery (Hafner et al., 2019). With respect to Alzheimer’s disease, this concept of accelerated translation by amyloid and tau causing a global break on translation has been shown elsewhere (Cruz et al., 2023; Li & Götz, 2017).

The global translation was significantly decreased in all mechanosensory tau strains suggesting tau’s involvement with other tissues to cause a systemic effect (**Fig. 2B**). We believe this could be due to the cell non-autonomous action of tau influencing translation in other tissues. It’s known that organismal proteostasis is governed by cell non-autonomous regulation in *C. elegans* especially regarding heat shock (Morimoto, 2020). Therefore, tau expressed in mechanosensory neurons could be affecting other tissues to cause global changes in translation. Our pan-neuronal strain didn’t show any significant impairment in lifespan or locomotion but global translation was significantly decreased (**Fig. 2D**). We attribute the neuronal translation effect to the presence of specific PTMs in this case T231 phosphorylation in mechanosensory tau strains. Considering the pro-longevity role of decreased translation (Rollins et al., 2019), we interpreted the results that tau wouldn’t be triggering pro-longevity related pathways because there’s still a significant decrease in lifespan (**Fig. 1A**).

We initially hypothesized that to cause ribosomal dysfunction, tau associates with actively translating ribosomes (polysomes). From our repeated attempts with fluorescent polysome profiling to resolve this association, we concluded that there’s no direct association between human tau and the ribosome (**Fig. 3A-E**). Throughout the profiling experiments, we altered our protocol with an addition of a fixating agent assuming a weak interaction and this also resulted in no association. We tried an *in vitro* strategy in which we acquired a human recombinant 0N4R tau protein and conjugate it to a fluorescent dye to be mixed with N2 lysate to saturate all ribosomes. Despite micrograms of tau protein mixed with the lysate, there was no association (**Fig. 4**). We also tried an *ex vivo* strategy in which we acquired frozen mouse cerebella where EGFP tagged tau is expressed pan-neuronally and to resolve the association with our standard protocol. This also resulted in no association. With our *in vitro*, *in vivo* and *ex vivo* approaches into fluorescent polysome profiling, we are confident that there is no direct association between tau and ribosome. Therefore, it is tempting to look for causal mechanisms to explain our neuronal and global translation phenotype. One of the strong contenders is a possible cell non-autonomous action of tau to influence translation levels in other tissues. Cell non-autonomous signaling is a very common phenomenon in *C. elegans* (Taylor et al., 2014). There are reports showing cell non-autonomous action from body wall muscle protein influencing neuronal guidance and intestinal proteins influencing neuropeptide signaling (Ochs et al., 2020; Savini et al., 2022). Tau expressed either in mechanosensory neurons or all neurons might trigger nearby tissues to decrease translation via an unknown mechanism.

There is another possibility of tau disrupting ribosome biogenesis in nucleolus (**Fig. 4E-F**). There is existing evidence of nuclear tau and the alleged role of tau in nucleus/nucleolus (Bukar Maina et al., 2016; Loomis et al., 1990; Maina, Bailey, Doherty, et al., 2018). In our imaging dataset, we did notice a nucleolar tau signal in mechanosensory tau strains (**Fig. 4E**). In an FTD mouse model, tau is known to decrease 60S large subunit synthesis and synthesis of ribosomal proteins (Evans et al., 2021; Nyhus et al., 2019). It is also known that even minor perturbations to ribosomal biogenesis could result in intellectual disability or microcephaly (Ni & Buszczak, 2022). Tau could be altering rRNA production and therefore ribosome biogenesis to cause global changes in translation (Antón-Fernández et al., 2023).

These findings might have important implications for the development of diagnostic and therapeutic strategies for AD. By identifying tau-ribosome association as a potential early event of disease pathology, it may be possible to intervene AD at an earlier stage before irreversible damage to the brain has occurred. Moreover, targeting tau-ribosome interactions may represent a novel therapeutic strategy for AD, aimed at restoring normal protein synthesis and neuronal function in the affected brain. In a mouse prion model, it was recently shown that by restoring global translation levels, neurodegeneration could be ameliorated (Albert-Gasco et al., 2024). Tau’s disruption of neuronal translation could also be the driving force for selective vulnerability (Kampmann, 2024).

In conclusion, the association between tau and the ribosome is an emerging area of research with important implications for our understanding of AD pathogenesis. Studies in *C. elegans* and other model systems have provided evidence that tau-ribosome association might represent an important early event in the disease process. We believe that tau is causing ribosomal dysfunction despite our results showing no association with the ribosome (**Fig. 2,3**). In addition, certain post-translational modifications of tau such as T231 phosphorylation and K274/281 acetylation might exacerbate ribosomal dysfunction. We also anticipate that *de novo* translation dependent mechanisms such as long-term associative learning could be affected by ribosomal function (Perez et al., 2021). Further research in this area is needed to elucidate the precise mechanisms by which wild type or modified tau disrupt ribosome function and to explore the potential of targeting ribosomal dysfunction for diagnostic and therapeutic purposes in tauopathies.

## Acknowledgements

We would like to acknowledge Dr. Frédéric X.A. Bonnet at the Light Microscopy Facility in MDI Biological Laboratory for his assistance with the imaging and OPP dataset. We would like to acknowledge Dr. Romain Madelaine at MDIBL for his valuable scientific input. We would like to thank Dr. Keith Nehrke at University of Rochester for the mechanosensory tau strains. We would like to thank Dr. Brian Ackley at University of Kansas for pan-neuronal tau strains. We would like to thank Dr. George Canet and Dr. Emmanuel Planel at Université Laval for the frozen mouse cerebellar tissue. We would like to acknowledge our funding sources Comparative Biology of Tissue, Repair, Regeneration and Aging (COBRE) grants P20GM104318-05, P20GM103423 from National Institute of General Medical Sciences (NIGMS) to J.A.R.

## References

Albert-Gasco, H., Smith, H. L., Alvarez-Castelao, B., Swinden, D., Halliday, M., Janaki-Raman, S., Butcher, A. J., & Mallucci, G. R. (2024). Trazodone rescues dysregulated synaptic and mitochondrial nascent proteomes in prion neurodegeneration. Brain, 147(2), 649–664. 10.1093/brain/awad313

Alquezar, C., Arya, S., & Kao, A. W. (2021). Tau Post-translational Modifications: Dynamic Transformers of Tau Function, Degradation, and Aggregation. Frontiers in Neurology, 11, 595532. 10.3389/fneur.2020.595532

Antón-Fernández, A., Vallés-Saiz, L., Avila, J., & Hernández, F. (2023). Neuronal nuclear tau and neurodegeneration. Neuroscience, 518, 178–184. 10.1016/j.neuroscience.2022.07.015

Aquino Nunez, W., Combs, B., Gamblin, T. C., & Ackley, B. D. (2022). Age-dependent accumulation of tau aggregation in Caenorhabditis elegans. Frontiers in Aging, 3, 928574. 10.3389/fragi.2022.928574

Banerjee, S., Ferdosh, S., Ghosh, A. N., & Barat, C. (2020). Tau protein-induced sequestration of the eukaryotic ribosome: Implications in neurodegenerative disease. Scientific Reports, 10(1), 5225. 10.1038/s41598-020-61777-7

Ben-Zvi, A., Miller, E. A., & Morimoto, R. I. (2009). Collapse of proteostasis represents an early molecular event in *Caenorhabditis elegans* aging. Proceedings of the National Academy of Sciences, 106(35), 14914–14919. 10.1073/pnas.0902882106

Boyarko, B., & Hook, V. (2021). Human Tau Isoforms and Proteolysis for Production of Toxic Tau Fragments in Neurodegeneration. Frontiers in Neuroscience, 15, 702788. 10.3389/fnins.2021.702788

Bukar Maina, M., Al-Hilaly, Y., & Serpell, L. (2016). Nuclear Tau and Its Potential Role in Alzheimer’s Disease. Biomolecules, 6(1), 9. 10.3390/biom6010009

Carroll, T., Guha, S., Nehrke, K., & Johnson, G. V. W. (2021). Tau Post-Translational Modifications: Potentiators of Selective Vulnerability in Sporadic Alzheimer’s Disease. Biology, 10(10), 1047. 10.3390/biology10101047

Cruz, E., Nisbet, R. M., & Götz, J. (2023). Break and accelerator—The mechanics of Tau (and amyloid) toxicity. Cytoskeleton, cm.21781. 10.1002/cm.21781

Dujardin, S., Commins, C., Lathuiliere, A., Beerepoot, P., Fernandes, A. R., Kamath, T. V., De Los Santos, M. B., Klickstein, N., Corjuc, D. L., Corjuc, B. T., Dooley, P. M., Viode, A., Oakley, D. H., Moore, B. D., Mullin, K., Jean-Gilles, D., Clark, R., Atchison, K., Moore, R., … Hyman, B. T. (2020). Tau molecular diversity contributes to clinical heterogeneity in Alzheimer’s disease. Nature Medicine, 26(8), 1256– 1263. 10.1038/s41591-020-0938-9

Evans, H. T., Taylor, D., Kneynsberg, A., Bodea, L.-G., & Götz, J. (2021). Altered ribosomal function and protein synthesis caused by tau. Acta Neuropathologica Communications, 9(1), 110. 10.1186/s40478-021-01208-4

Frontzkowski, L., Ewers, M., Brendel, M., Biel, D., Ossenkoppele, R., Hager, P., Steward, A., Dewenter, A., Römer, S., Rubinski, A., Buerger, K., Janowitz, D., Binette, A. P., Smith, R., Strandberg, O., Carlgren, N. M., Dichgans, M., Hansson, O., & Franzmeier, N. (2022). Earlier Alzheimer’s disease onset is associated with tau pathology in brain hub regions and facilitated tau spreading. Nature Communications, 13(1), 4899. 10.1038/s41467-022-32592-7

Fusco, C. M., Desch, K., Dörrbaum, A. R., Wang, M., Staab, A., Chan, I. C. W., Vail, E., Villeri, V., Langer, J. D., & Schuman, E. M. (2021). Neuronal ribosomes exhibit dynamic and context-dependent exchange of ribosomal proteins. Nature Communications, 12(1), 6127. 10.1038/s41467-021-26365-x

Gamarra, M., de la Cruz, A., Blanco-Urrejola, M., & Baleriola, J. (2021). Local Translation in Nervous System Pathologies. Frontiers in Integrative Neuroscience, 15, 689208. 10.3389/fnint.2021.689208

Guha, S., Fischer, S., Johnson, G. V. W., & Nehrke, K. (2020). Tauopathy-associated tau modifications selectively impact neurodegeneration and mitophagy in a novel C. elegans single-copy transgenic model. Molecular Neurodegeneration, 15(1), 65. 10.1186/s13024-020-00410-7

Guisle, I., Canet, G., Pétry, S., Fereydouni-Forouzandeh, P., Morin, F., Kérauden, R., Whittington, R. A., Calon, F., Hébert, S. S., & Planel, E. (2022). Sauna-like conditions or menthol treatment reduce tau phosphorylation through mild hyperthermia. Neurobiology of Aging, 113, 118–130. 10.1016/j.neurobiolaging.2022.02.011

Gunawardana, C. G., Mehrabian, M., Wang, X., Mueller, I., Lubambo, I. B., Jonkman, J. E. N., Wang, H., & Schmitt-Ulms, G. (2015). The Human Tau Interactome: Binding to the Ribonucleoproteome, and Impaired Binding of the Proline-to-Leucine Mutant at Position 301 (P301L) to Chaperones and the Proteasome. Molecular & Cellular Proteomics, 14(11), 3000–3014. 10.1074/mcp.M115.050724

Guo, T., Zhang, D., Zeng, Y., Huang, T. Y., Xu, H., & Zhao, Y. (2020). Molecular and cellular mechanisms underlying the pathogenesis of Alzheimer’s disease. Molecular Neurodegeneration, 15(1), 40. 10.1186/s13024-020-00391-7

Hacisuleyman, E., Hale, C. R., Noble, N., Luo, J., Fak, J. J., Saito, M., Chen, J., Weissman, J. S., & Darnell, R. B. (2024). Neuronal activity rapidly reprograms dendritic translation via eIF4G2:uORF binding. Nature Neuroscience, 27(5), 822–835. 10.1038/s41593-024-01615-5

Hafner, A.-S., Donlin-Asp, P. G., Leitch, B., Herzog, E., & Schuman, E. M. (2019). Local protein synthesis is a ubiquitous feature of neuronal pre- and postsynaptic compartments. Science, 364(6441), eaau3644. 10.1126/science.aau3644

Ittner, L. M., Ke, Y. D., Delerue, F., Bi, M., Gladbach, A., van Eersel, J., Wölfing, H., Chieng, B. C., Christie, M. J., Napier, I. A., Eckert, A., Staufenbiel, M., Hardeman, E., & Götz, J. (2010). Dendritic Function of Tau Mediates Amyloid-β Toxicity in Alzheimer’s Disease Mouse Models. Cell, 142(3), 387–397. 10.1016/j.cell.2010.06.036

Kampmann, M. (2024). Molecular and cellular mechanisms of selective vulnerability in neurodegenerative diseases. Nature Reviews Neuroscience, 25(5), 351–371. 10.1038/s41583-024-00806-0

Kavanagh, T., Halder, A., & Drummond, E. (2022). Tau interactome and RNA binding proteins in neurodegenerative diseases. Molecular Neurodegeneration, 17(1), 66. 10.1186/s13024-022-00572-6

Koren, S. A., Hamm, M. J., Meier, S. E., Weiss, B. E., Nation, G. K., Chishti, E. A., Arango, J. P., Chen, J., Zhu, H., Blalock, E. M., & Abisambra, J. F. (2019). Tau drives translational selectivity by interacting with ribosomal proteins. Acta Neuropathologica, 137(4), 571–583. 10.1007/s00401-019-01970-9

Li, C., & Götz, J. (2017). Somatodendritic accumulation of Tau in Alzheimer’s disease is promoted by Fyn-mediated local protein translation. The EMBO Journal, 36(21), 3120–3138. 10.15252/embj.201797724

Long, J. M., & Holtzman, D. M. (2019). Alzheimer Disease: An Update on Pathobiology and Treatment Strategies. Cell, 179(2), 312–339. 10.1016/j.cell.2019.09.001

Loomis, P. A., Howard, T. H., Castleberry, R. P., & Binder, L. I. (1990). Identification of nuclear tau isoforms in human neuroblastoma cells. Proceedings of the National Academy of Sciences, 87(21), 8422– 8426. 10.1073/pnas.87.21.8422

Maina, M. B., Bailey, L. J., Doherty, A. J., & Serpell, L. C. (2018). The Involvement of Aβ42 and Tau in Nucleolar and Protein Synthesis Machinery Dysfunction. Frontiers in Cellular Neuroscience, 12, 220. 10.3389/fncel.2018.00220

Maina, M. B., Bailey, L. J., Wagih, S., Biasetti, L., Pollack, S. J., Quinn, J. P., Thorpe, J. R., Doherty, A. J., & Serpell, L. C. (2018). The involvement of tau in nucleolar transcription and the stress response. Acta Neuropathologica Communications, 6(1), 70. 10.1186/s40478-018-0565-6

Mandelkow, E.-M., & Mandelkow, E. (2012). Biochemistry and Cell Biology of Tau Protein in Neurofibrillary Degeneration. Cold Spring Harbor Perspectives in Medicine, 2(7), a006247–a006247. 10.1101/cshperspect.a006247

Marck, A., Berthelot, G., Foulonneau, V., Marc, A., Antero-Jacquemin, J., Noirez, P., Bronikowski, A. M., Morgan, T. J., Garland, T., Carter, P. A., Hersen, P., Di Meglio, J.-M., & Toussaint, J.-F. (2016). Age-Related Changes in Locomotor Performance Reveal a Similar Pattern for *Caenorhabditis elegans*, *Mus domesticus*, *Canis familiaris*, *Equus caballus*, and *Homo sapiens*. The Journals of Gerontology Series A: Biological Sciences and Medical Sciences, glw136. 10.1093/gerona/glw136

Meier, S., Bell, M., Lyons, D. N., Rodriguez-Rivera, J., Ingram, A., Fontaine, S. N., Mechas, E., Chen, J., Wolozin, B., LeVine, H., Zhu, H., & Abisambra, J. F. (2016). Pathological Tau Promotes Neuronal Damage by Impairing Ribosomal Function and Decreasing Protein Synthesis. Journal of Neuroscience, 36(3), 1001–1007. 10.1523/JNEUROSCI.3029-15.2016

Monday, H. R., Kharod, S. C., Yoon, Y. J., Singer, R. H., & Castillo, P. E. (2022). Presynaptic FMRP and local protein synthesis support structural and functional plasticity of glutamatergic axon terminals. Neuron, 110(16), 2588–2606.e6. 10.1016/j.neuron.2022.05.024

Morimoto, R. I. (2020). Cell-Nonautonomous Regulation of Proteostasis in Aging and Disease. Cold Spring Harbor Perspectives in Biology, 12(4), a034074. 10.1101/cshperspect.a034074

Natale, C., Barzago, M. M., & Diomede, L. (2020). Caenorhabditis elegans Models to Investigate the Mechanisms Underlying Tau Toxicity in Tauopathies. Brain Sciences, 10(11), 838. 10.3390/brainsci10110838

Ni, C., & Buszczak, M. (2022). The homeostatic regulation of ribosome biogenesis. *Seminars in Cell & Developmental Biology*, S1084952122001252. 10.1016/j.semcdb.2022.03.043

Nyhus, C., Pihl, M., Hyttel, P., & Hall, V. J. (2019). Evidence for nucleolar dysfunction in Alzheimer’s disease. Reviews in the Neurosciences, 30(7), 685–700. 10.1515/revneuro-2018-0104

Ochs, M. E., Josephson, M. P., & Lundquist, E. A. (2020). The Predicted RNA-Binding Protein ETR-1/CELF1 Acts in Muscles To Regulate Neuroblast Migration in *Caenorhabditis elegans*. G3 Genes|Genomes|Genetics, 10(7), 2365–2376. 10.1534/g3.120.401182

Ostroff, L. E., & Cain, C. K. (2022). Persistent up-regulation of polyribosomes at synapses during long-term memory, reconsolidation, and extinction of associative memory. Learning & Memory (Cold Spring Harbor, N.Y.), 29(8), 192–202. 10.1101/lm.053577.122

Perez, J. D., Fusco, C. M., & Schuman, E. M. (2021). A Functional Dissection of the mRNA and Locally Synthesized Protein Population in Neuronal Dendrites and Axons. Annual Review of Genetics, 55(1), 183–207. 10.1146/annurev-genet-030321-054851

Pringle, E. S., McCormick, C., & Cheng, Z. (2019). Polysome Profiling Analysis of mRNA and Associated Proteins Engaged in Translation. Current Protocols in Molecular Biology, 125(1), e79. 10.1002/cpmb.79

Raveh, A., Margaliot, M., Sontag, E. D., & Tuller, T. (2016). A model for competition for ribosomes in the cell. Journal of The Royal Society Interface, 13(116), 20151062. 10.1098/rsif.2015.1062

Rollins, J. A., Shaffer, D., Snow, S. S., Kapahi, P., & Rogers, A. N. (2019). Dietary restriction induces posttranscriptional regulation of longevity genes. Life Science Alliance, 2(4), e201800281. 10.26508/lsa.201800281

Savini, M., Folick, A., Lee, Y.-T., Jin, F., Cuevas, A., Tillman, M. C., Duffy, J. D., Zhao, Q., Neve, I. A., Hu, P.-W., Yu, Y., Zhang, Q., Ye, Y., Mair, W. B., Wang, J., Han, L., Ortlund, E. A., & Wang, M. C. (2022). Lysosome lipid signalling from the periphery to neurons regulates longevity. Nature Cell Biology. 10.1038/s41556-022-00926-8

Schwalbe, M., Kadavath, H., Biernat, J., Ozenne, V., Blackledge, M., Mandelkow, E., & Zweckstetter, M. (2015). Structural Impact of Tau Phosphorylation at Threonine 231. Structure, 23(8), 1448–1458. 10.1016/j.str.2015.06.002

Shaffer, D., & Rollins, J. (2020). Fluorescent Polysome Profiling in Caenorhabditis elegans. BIO-PROTOCOL, 10(17). 10.21769/BioProtoc.3742

Sohn, P. D., Tracy, T. E., Son, H.-I., Zhou, Y., Leite, R. E. P., Miller, B. L., Seeley, W. W., Grinberg, L. T., & Gan, L. (2016). Acetylated tau destabilizes the cytoskeleton in the axon initial segment and is mislocalized to the somatodendritic compartment. Molecular Neurodegeneration, 11(1), 47. 10.1186/s13024-016-0109-0

Somers, H. M., Fuqua, J. H., Bonnet, F. X. A., & Rollins, J. A. (2022). Quantification of tissue-specific protein translation in whole C. elegans using O-propargyl-puromycin labeling and fluorescence microscopy. Cell Reports Methods, 2(4), 100203. 10.1016/j.crmeth.2022.100203

Sun, C., Nold, A., Fusco, C. M., Rangaraju, V., Tchumatchenko, T., Heilemann, M., & Schuman, E. M. (2021). The prevalence and specificity of local protein synthesis during neuronal synaptic plasticity. Science Advances, 7(38), eabj0790. 10.1126/sciadv.abj0790

Taylor, R. C., Berendzen, K. M., & Dillin, A. (2014). Systemic stress signalling: Understanding the cell non-autonomous control of proteostasis. Nature Reviews Molecular Cell Biology, 15(3), 211–217. 10.1038/nrm3752

Tracy, T. E., Madero-Pérez, J., Swaney, D. L., Chang, T. S., Moritz, M., Konrad, C., Ward, M. E., Stevenson, E., Hüttenhain, R., Kauwe, G., Mercedes, M., Sweetland-Martin, L., Chen, X., Mok, S.-A., Wong, M. Y., Telpoukhovskaia, M., Min, S.-W., Wang, C., Sohn, P. D., … Gan, L. (2022). Tau interactome maps synaptic and mitochondrial processes associated with neurodegeneration. Cell, 185(4), 712–728.e14. 10.1016/j.cell.2021.12.041

Tracy, T. E., Sohn, P. D., Minami, S. S., Wang, C., Min, S.-W., Li, Y., Zhou, Y., Le, D., Lo, I., Ponnusamy, R., Cong, X., Schilling, B., Ellerby, L. M., Huganir, R. L., & Gan, L. (2016). Acetylated Tau Obstructs KIBRA-Mediated Signaling in Synaptic Plasticity and Promotes Tauopathy-Related Memory Loss. Neuron, 90(2), 245–260. 10.1016/j.neuron.2016.03.005

Wang, X., Williams, D., Müller, I., Lemieux, M., Dukart, R., Maia, I. B. L., Wang, H., Woerman, A. L., & Schmitt-Ulms, G. (2019). Tau interactome analyses in CRISPR-Cas9 engineered neuronal cells reveal ATPase-dependent binding of wild-type but not P301L Tau to non-muscle myosins. Scientific Reports, 9(1), 16238. 10.1038/s41598-019-52543-5

Wesseling, H., Mair, W., Kumar, M., Schlaffner, C. N., Tang, S., Beerepoot, P., Fatou, B., Guise, A. J., Cheng, L., Takeda, S., Muntel, J., Rotunno, M. S., Dujardin, S., Davies, P., Kosik, K. S., Miller, B. L., Berretta, S., Hedreen, J. C., Grinberg, L. T., … Steen, J. A. (2020). Tau PTM Profiles Identify Patient Heterogeneity and Stages of Alzheimer’s Disease. Cell, 183(6), 1699–1713.e13. 10.1016/j.cell.2020.10.029

